# Temporal Hydrogen-Bond Network Analysis Reveals Substrate-Directed Connectivity in Dihydrofolate Reductase

**DOI:** 10.64898/2026.05.05.722848

**Authors:** Tandac F. Guclu, Canan Atilgan, Ali Rana Atilgan

## Abstract

Hydrogen-bond networks are central to protein function, but most network analyses rely on static representations that neglect how interactions evolve in time. Here, we introduce a framework that combines instantaneous and temporal graph analysis of hydrogen-bond networks derived from molecular dynamics trajectories to quantify ligand-directed hydrogen-bond connectivity. We apply the method to *E. coli* dihydrofolate reductase (DHFR) and its L28R mutant, computing shortest hydrogen-bond paths from all residues to the substrate dihydrofolate (DHF). The instantaneous analysis reveals that DHF-directed connectivity is organized through a sparse set of preferred routes, with D27 and T113 acting as prominent hubs in the wild-type enzyme. Temporal analysis highlights residues that preferentially support time-ordered DHF-directed connectivity. Comparison with L28R shows that the mutation preserves the main substrate-contacting architecture and the overall communication scaffold but redistributes pathway usage, persistence, and temporal convergence. The network-derived hotspots partially overlap with independent coevolution signals, most strongly in the K109–I115 region, while overlap with cryptic-site predictors is more limited. This pattern indicates that the hydrogen-bond network captures evolutionarily supported communication regions in DHFR that are not fully recovered by static structural approaches. The framework is broadly applicable to ligand-binding proteins and provides a route to identify persistent, delayed, and mutation-sensitive signaling pathways directly from time-ordered simulation data.

## Introduction

The wide range of protein functions, from enzymatic catalysis to signal transduction, is organized by the dynamic nature of internal interaction networks within protein structures [1–11]. Hydrogen bonds are particularly well suited for describing these networks because they are directional, reversible, and sensitive to environmental and sequence changes [3, 8, 10–14]. Hydrogen-bonding patterns therefore provide a practical basis for interrogating how mutations reshape connectivity and how residues that do not form direct contacts remain functionally coupled, as in allosteric regulation [10, 15–17]. Distal effects are also relevant for inhibition because proteins can harbor cryptic sites. These are transient pockets away from the functional site that become druggable only in specific conformations [18–25]. Cryptic sites are commonly assessed by analyzing the protein surface and evaluating whether an emerging cavity resembles a native binding site [21, 22, 24, 26]. In addition, coevolution can highlight residues that are structurally and functionally coupled even when they are distant in sequence [27–31]. All these observations motivate methods that can identify indirect residue–residue coupling in a manner consistent with the underlying conformational dynamics.

Allosteric communication is a hierarchical, multi-timescale phenomenon in which nanosecond-scale motions act as the earliest carriers of long-range signals. Transient infrared spectroscopy of photo-switchable PDZ domains has shown that ligand-induced allosteric transitions are initiated within nanoseconds and propagate through population shifts among metastable states rather than through large mean structural changes [32, 33]. Similar nanosecond-to-microsecond hierarchies have been resolved in other allosteric and photosensor systems [34–36], establishing nanosecond motions as a fundamental layer of allosteric communication that is directly accessible to molecular dynamics-based hydrogen-bond network analysis. At the same time, full propagation of allosteric signals can extend into longer timescales not always covered by current trajectory lengths.

Molecular dynamics (MD) simulations provide direct access to time-dependent conformations, but extracting temporally ordered connectivity pathways from trajectories remains challenging. Graph-based representations offer a coarsened description in which residues or atoms define nodes and interactions define edges, enabling the use of metrics such as shortest-path length, betweenness centrality, and clustering coefficients to identify communication hubs and quantify mutational impacts on long-range coupling [7, 8, 37–39]. However, most analyses rely on static graphs constructed from average structures or independent snapshots [40–42]. Such representations implicitly treat all edges as simultaneously available and do not capture the temporal ordering of interaction formation and disruption. For hydrogen-bond networks, this limitation is particularly consequential because a path is intrinsically time dependent [43]. It has a lifetime over which it remains connected and a latency that reflects the time required for connectivity to propagate from a source to a sink. Methods that explicitly incorporate this temporal causality are therefore needed to connect instantaneous connectivity to dynamic communication.

Here we introduce a computational framework that integrates classical hydrogen-bond analysis with graph-based and temporal network descriptions to quantify ligand-directed connectivity in proteins. From atomistic MD trajectories, we construct an instantaneous hydrogen-bond network at each snapshot and compute shortest paths from all residues to a chosen molecular target, also explicitly considering water molecules [3, 4, 10, 13, 14, 17, 44]. This network view yields path-length distributions and identifies source residues that preferentially feed paths into the ligand. In parallel, we formulate a temporal network description in which hydrogen bonds and their paths are treated as events with defined start and end times. By tracking consecutive occurrences of bonds and paths, we quantify their respective lifetimes, and path completion times that report how efficiently connectivity can propagate when edges must exist in sequence. The combined analysis provides complementary static and temporal descriptors derived from the same trajectory.

We demonstrate the approach using *E. coli* dihydrofolate reductase (DHFR), a well-established model enzyme whose catalytic cycle couples ligand binding to conformational changes and long-range communication [8, 9, 45–50]. DHFR mutations have also been studied experimentally, and substitutions can modulate binding affinity and kinetics through subtle changes rather than large structural rearrangements, making it a useful test case for path-based analyses. We apply the framework to wild-type (WT) DHFR and to the L28R mutant, an extensively studied variant [8, 9, 46, 51–53], using this targeted perturbation to evaluate how a single amino acid change reshapes substrate-directed hydrogen-bond connectivity, path persistence, and temporal accessibility. The procedure relies only on hydrogen-bond geometries and time-ordered trajectories, making it readily transferable to other protein systems.

## Materials & Methods

### Simulations of wild-type and L28R mutant

The 3QL3 structure was downloaded from the Protein Data Bank [54] as the reference WT structure. The system consists of the protein DHFR, the cofactor nicotinamide adenine dinucleotide phosphate (NADPH), and the substrate dihydrofolate (DHF).

We then used the CHARMM36 [55] force field; the parameters for NADPH and DHF are the same as those used in our previous study [8]. The system was solvated in a rectangular water box with a minimum distance of 10 Å in all directions. KCl was added to neutralize the system and reach an isotonic salt concentration of 0.15 M. MD simulations were performed using NAMD3 [56] at 310 K and 1 atm under constant temperature and pressure conditions. Each simulation was run for 1 µs and replicated once, for a total of 2 µs.

The mutated system (L28R) was generated using a free energy perturbation scheme [57], in which the hybrid amino acid side chain was tuned throughout the trajectory to obtain an equilibrated conformation. The final frame of this equilibration was then used to initiate two independent 1 µs simulations.

### Hydrogen-bond analyses

Hydrogen bonds were computed using the MDAnalysis [58] Python package, with the definition of a donor–acceptor distance ≤ 3.2 Å and a donor–hydrogen–acceptor angle > 150° (**Figure 1a**). We performed hydrogen-bond analysis on the DHFR protein, NADPH, DHF, and on vicinal water molecules [59] located within 4 Å of the protein and the ligands. All possible hydrogen bonds were computed between protein-system residues (including the ligands), between the protein system and water, and between these vicinal water molecules. The time evolution of hydrogen bonds was saved for further residue graph construction. The 1 ns sampling interval used here matches the nanosecond timescale that has been independently identified as the earliest carrier of allosteric signal propagation in time-resolved IR studies of small allosteric domains [32].

**Figure 1.**
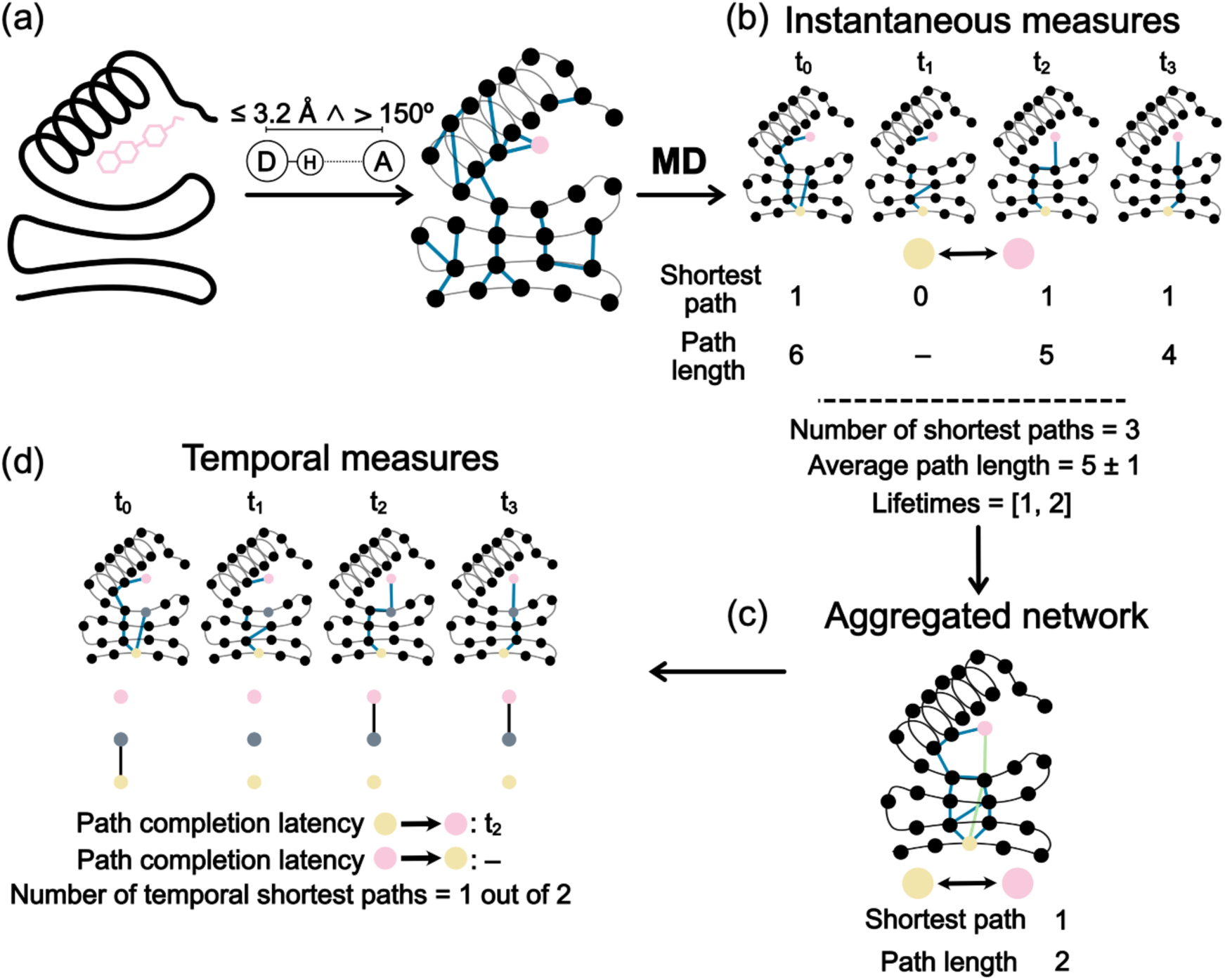
Overview of the hydrogen-bond network workflow used to extract instantaneous and temporal paths from MD trajectories. **(a)** Hydrogen bonds were identified in each MD snapshot using geometric criteria (donor–acceptor distance ≤ 3.2 Å and donor–hydrogen–acceptor angle > 150°) and converted into residue-level interaction networks containing DHFR, NADPH, DHF, and vicinal water molecules. **(b)** Instantaneous measures were obtained by computing shortest paths independently in each snapshot, yielding snapshot-wise connectivity, path lengths, path counts, and path lifetimes from consecutive occurrences. A source node is exemplified in yellow; the target in this work is always the DHF node, shown in pink. Hydrogen bonds in each snapshot are shown in teal. **(c)** All nodes and edges observed across the trajectory were then merged into an aggregated network. The edges belonging to the shortest path in the aggregated graph are shown in green. **(d)** Temporal measures were obtained by testing whether shortest paths defined on the aggregated graph can be completed sequentially over time. In this example, only one of the two possible temporal paths is completed, illustrating the asymmetry of temporal shortest paths: because edges must be traversed in sequential time order, completion in one direction does not imply completion in the reverse direction. The gray node marks an arbitrary central node through which many temporal paths pass; such nodes are therefore expected to have a large temporal incoming path count.

### Residue network generation

Information from hydrogen bonds was used to generate protein graphs. We included all 159 residues of DHFR, NADPH and DHF, yielding a total of 161 nodes for the non-water part. In addition, vicinal water molecules were treated as nodes; their residue indices were recorded at every snapshot and labeled as the water component. Throughout the WT simulation, an average of 977 ± 124 vicinal water molecules were computed. The graphs are undirected and contain no self-loops. There are also no parallel edges, meaning that nodes connected by more than one hydrogen bond are represented by a single edge. Because the vicinal water environment is highly dynamic, the number of nodes comprising the solvent network varies throughout the simulation trajectory. To map this consistently, we record all unique molecular identities across the trajectories and construct a master adjacency matrix that captures their neighborhood connectivity [60, 61]. This approach yields a total of 1462 unique nodes, encompassing both protein residues and water molecules, for the WT simulations. By analyzing this overarching network, we track the changing edge patterns over time, observing that unconnected (dangling) nodes continuously appear as each snapshot forms an instantaneous residue graph of the protein and solvent components. By extracting snapshots at 1 ns intervals from the two 1 µs simulations, we generated a total of 2000 residue graphs for our analysis. Note that, because the networks are constructed over hydrogen bonds, they are not necessarily fully connected.

### Analyses of residue networks

All calculations were performed using the NetworkX [62] package. Shortest paths were computed using the breadth-first search algorithm. To assess the temporal persistence of connections, the consecutive occurrences of edges and paths were tracked and defined as their lifetimes (**Figure 1b**). We recorded the lengths of these lifetimes to compute their overall probability distributions. For instance, if the existence of an edge or path between two nodes across ten snapshots is given by the array [1, 0, 0, 0, 1, 1, 1, 0, 1, 1], the resulting lifetimes are [1, 3, 2], which are determined by counting the continuous sequences of ones (**Figure 1b**).

We hypothesize that paths connecting to the substrate DHF are crucial for binding and may also reflect distal regulation (allostery). Hence, we focused on paths that end at DHF along the trajectory. For each of the instantaneous graphs generated previously, we computed all shortest paths from every residue to DHF. We refer to the analyses performed on these individual snapshots as instantaneous measures (**Figure 1b**). The shortest paths were classified based on whether they included water molecules as nodes. We computed path lengths to quantify the number of edges required for connectivity between each residue and DHF (shown respectively in yellow and pink in **Figure 1b**). After analyzing these lengths per snapshot, we calculated the time-averaged path length for each residue. In addition, we calculated the lifetimes of paths originating from each residue and time-averaged them to assess the persistence of recurring connections.

To quantify the structural connectivity converging on a specific intermediate residue, *x*, within the residue network, we calculated the Incoming Path Count. This metric evaluates the total number of shortest pathways originating from all other residues in the protein that pass through a given intermediate residue to reach DHF. Initially, this is conceptualized as the total number of shortest paths from any source residue to DHF that traverse the intermediate residue, divided by the number of shortest paths connecting the intermediate residue to DHF:

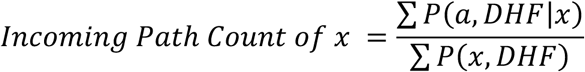

where ∑ *P*(*a*, *DHF*|*x*) indicates the total number of shortest paths starting from node *a* and ending at DHF that traverse through node *x*. ∑ *P*(*x*, *DHF*) represents the number of shortest paths between nodes *x* and DHF. This calculation can be simplified by using Bellman’s principle of optimality [63]. Assuming these pathway segments combine independently, the total number of such paths is the product of the two segments:

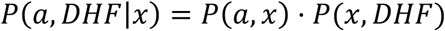

*P*(*x*, *DHF*) cancels out and leaves *P*(*a*, *x*), where the number of all shortest paths that lead to node *x* is approximated by, *Incoming Path Count of x* = ∑ *P*(*a*, *x*)

∑ *P*(*a*, *DHF*|*x*) is highly correlated with the time-averaged DHF-directed betweenness centrality computed on the instantaneous graphs (**Figure S1**, *R*^2^ = 0.98). Previously, we used betweenness centrality [64] to investigate protein function and to identify residues that dominate inter-residue connectivity [8, 37]. In the present work, however, we use the Incoming Path Count because it provides a simpler and more computationally efficient description of substrate-directed connectivity. Both measures quantify how strongly a residue contributes to shortest paths toward DHF, but the Incoming Path Count simplifies the calculation by collapsing the downstream branching from *a* to DHF and retaining only the accumulated inflow of source-to-DHF shortest paths through *a*. In the context of protein dynamics, this simplification highlights residues that act as funnels or integrators of DHF-directed connectivity. The strong agreement between ∑ *P*(*a*, *DHF*|*x*) and betweenness centrality arises from the sparse nature of the instantaneous hydrogen-bond graphs, in which most connected node pairs are linked by a unique shortest path (see Results below); under these conditions, the normalization that distinguishes betweenness centrality is usually trivial, and the two measures become nearly equivalent in practice. This simplified formulation is especially advantageous for the temporal analysis, where path definitions are ordered, asymmetric, and latency-dependent, making a full betweenness-style treatment less tractable and less transparent algorithmically.

Since ligand-induced allosteric transmission through the protein structure can occur with a time lag, we examined whether hydrogen-bond paths complete traversal from residues to DHF not within the same snapshot, but over the trajectory after some elapsed time. To this end, we first generated an aggregated graph (**Figure 1c**), in which we included all nodes (161 nodes from the protein and all water molecules that ever enter the vicinal region) and tracked the completion of these paths over the trajectory (**Figure 1d**). We recorded the completion time to assess how fast these paths were completed, because, compared to the instantaneous graphs, paths derived from the aggregated graph are spatially shorter. In this view, we assume that there is a waiting time on the nodes along the aggregated shortest paths and we track the time to reach the sink through these nodes. We refer to these analyses as temporal measures (**Figure 1d**). These measures are asymmetric; therefore, calculations were performed in both directions (from residues to DHF and from DHF to residues). For each direction, the path completion time was calculated for all shortest paths between the source and the sink and then averaged. Furthermore, the Incoming Path Count measure was also applied to temporal shortest paths to identify residues that control pathways with latency.

The pipeline described above operates on hydrogen-bond analyses applied to time-ordered MD trajectories and is therefore not specific to DHFR. The same procedure can be applied to any protein, either with a target ligand or molecule as the sink, or between arbitrary residue–residue pairs, with the set of nodes tailored to the system of interest, after which the instantaneous and temporal descriptors are computed in the same way. A reference Python implementation is provided on both GitHub and Zenodo.

## Results & Discussion

### Hydrogen-bond network and baseline statistics of the WT system

The residue networks of WT DHFR are constructed from 159 protein residues, NADPH, DHF, and vicinal water molecules, yielding 1462 unique nodes across the trajectory. The number of edges fluctuates around 200 throughout the two 1 µs simulations (**Figure S2**). The three classes of hydrogen bonds exhibit the expected hierarchy of lifetimes (**Figure S3**): protein–protein bonds are the longest-lived, protein–water bonds are intermediate, and water–water bonds are the shortest-lived. Because the vicinal water environment reorients on a sub-nanosecond timescale, our 1 ns sampling interval captures water–water interactions predominantly as single-frame events, consistent with prior reports [13, 16, 43, 65] and motivating our explicit treatment of temporal effects.

Within the instantaneous networks, connected node pairs are joined by a single shortest path in the overwhelming majority of snapshots (**Figure S4a**). Shortest-path lengths are sharply distributed, with direct contacts (length 1) most frequent and a gradually decaying tail extending up to 15 edges in rare snapshots (**Figure S4b**). Consecutive water molecules within paths are typically limited to one to four, with extended bridges of up to 13 waters occurring sporadically (**Figure S4c**). This sparse, short-path topology defines the baseline on which the DHF-directed analyses below are built.

### Instantaneous DHF-directed paths reveal a sparse communication scaffold in WT DHFR

DHFR function depends on its interactions with the substrate DHF (**Figure 2a**), and the hydrogen bonds that form the immediate binding interface are summarized in **Figure 2b,d**. Across 2000 snapshots, three residues dominate the direct-contact interface: D27, K32, and R52, with occurrences of 1708, 1140, and 1718, respectively. The remaining interface residues (R57, N23, A6, T35, S49, T113, N18) each form direct contacts in fewer than 250 snapshots and participate transiently.

**Figure 2.**
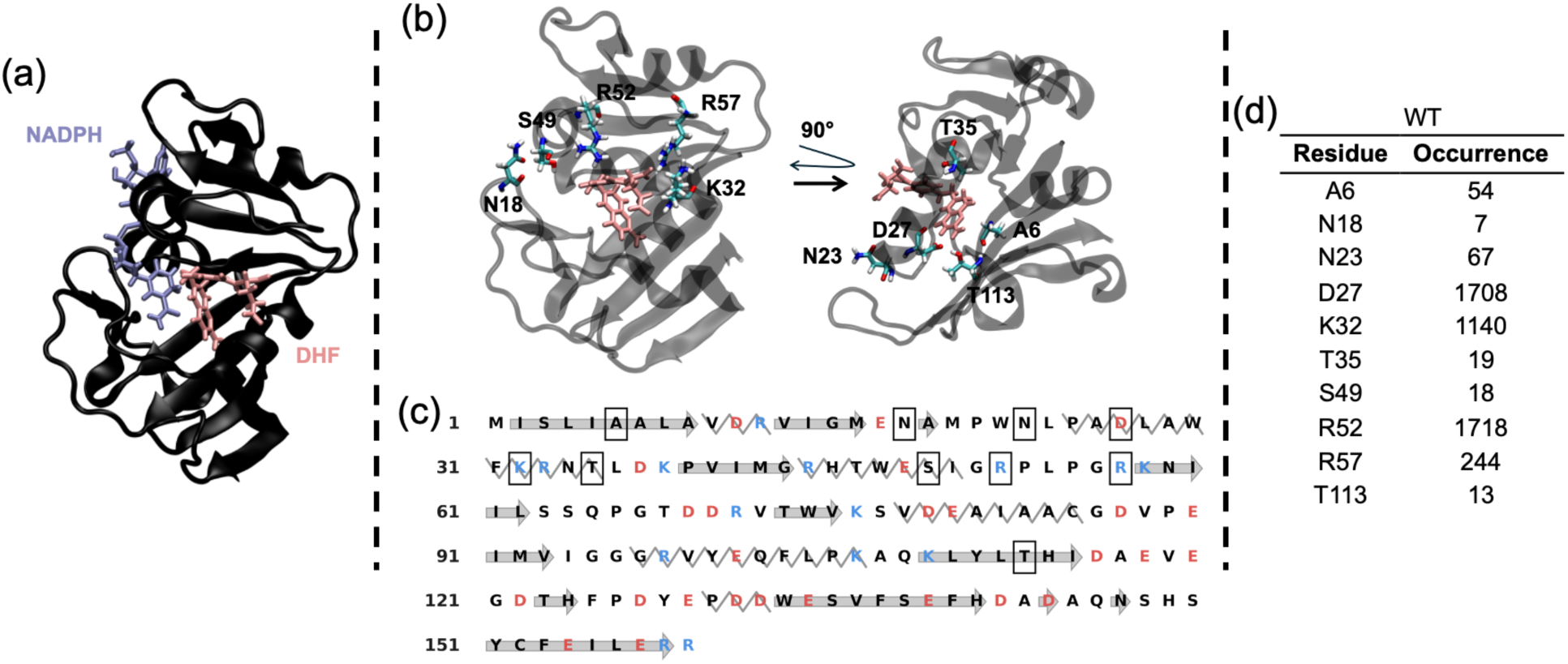
(a) The PDB structure of the simulated system, consisting of the DHFR protein, the cofactor NADPH, and the substrate DHF. Water molecules are omitted from these visualizations. **(b)** Close-up structural views highlighting specific amino acids in direct contact with DHF throughout the WT simulations. **(c)** The sequence map of the protein. Secondary structures are indicated with helices as zigzags and β-sheets as arrows. Residues making direct contact with DHF are boxed. Charged residues are color-coded: positive in blue and negative in red. **(d)** Occurrences of direct hydrogen bonds between WT DHFR residues and the DHF substrate. The occurrences represent the number of times a direct hydrogen bond was observed between the specified residue and DHF across 2000 snapshots extracted from two 1 µs molecular dynamics simulations. Residues D27, K32, and R52 exhibit the most frequent direct interactions.

To examine connectivity beyond the immediate interface, we computed all shortest paths from every residue to DHF across the 2000 instantaneous graphs, classifying each path as dry, wet, or total (**Figure 3a**). Five clusters of residues dominate the DHF-directed path count: (i) L4/A6/L8, (ii) L24/D27/F31/K32/T35/K38, (iii) T46/I50/R52/P55/R57, (iv) E90 as an isolated peak, and (v) K109/Y111/T113/I115. Structurally, these distal clusters do not reach the substrate through independent routes; rather, they funnel through a small set of shared intermediates (exemplified in **Figure 3b–d**). **Table 1** makes this explicit: the most populated recurring paths converge on the T113–D27 pair, with alternative branches through K32 and R52. Wet paths contribute only weakly to the total count, indicating that residue-mediated routes dominate snapshot-wise connectivity.

**Figure 3.**
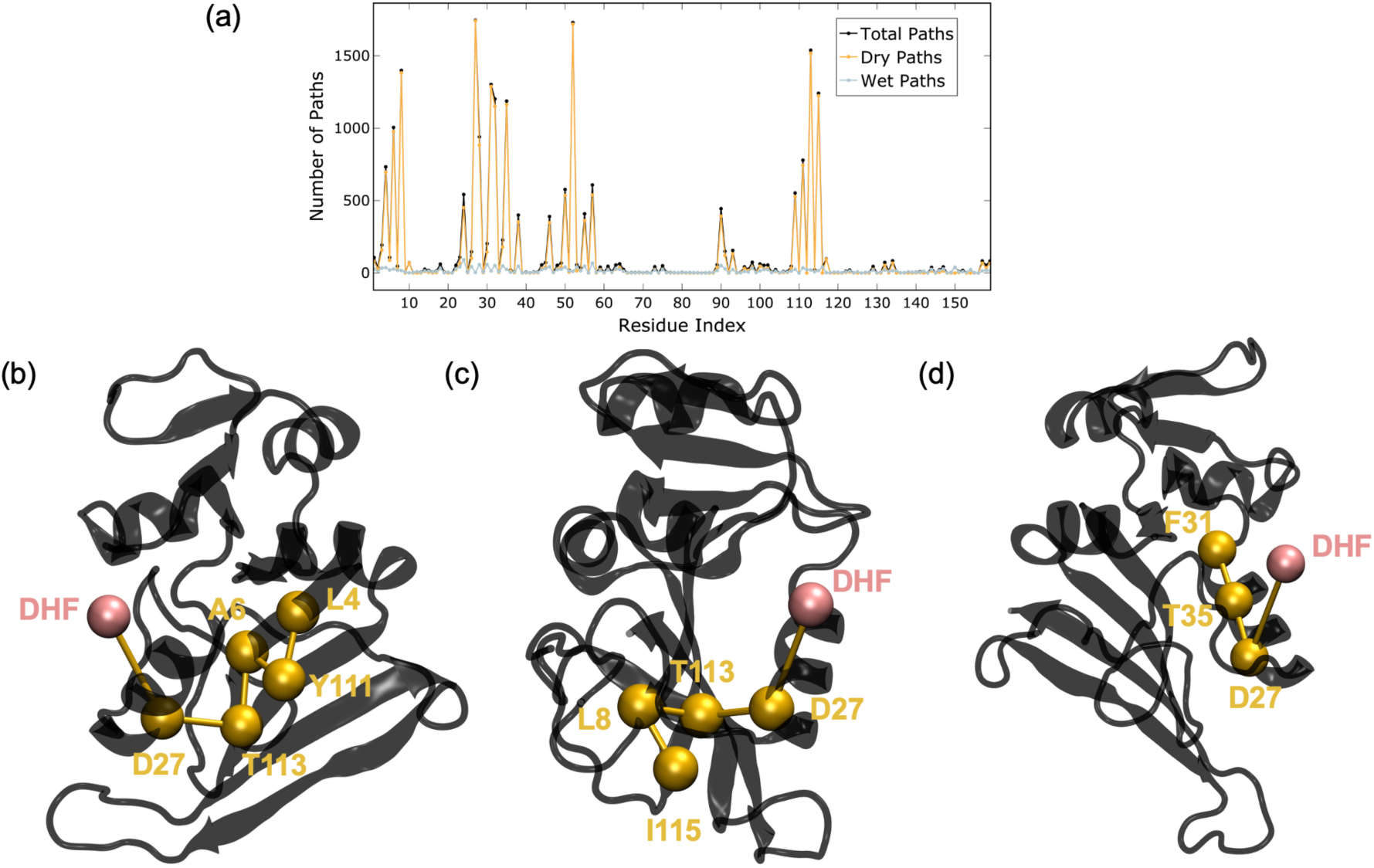
Hydrogen-bond shortest paths directed toward the DHF substrate in the WT system. **(a)** The total count of shortest paths originating from each residue and ending at DHF, computed across 2000 MD snapshots. The analysis categorizes pathways into dry paths (excluding water nodes), wet paths (those with water nodes), and total paths. Prominent peaks identify clusters of residues with frequent connectivity to DHF, including the L4/A6/L8 cluster, the L24–K38 region, the T46–R57 region, and the K109–I115 cluster. **(b–d)** Structural representations of highly recurring paths. These panels illustrate how shortest paths originating from distal residues funnel through key hub-like mediating residues such as D27 and T113 to reach the DHF substrate.

**Table 1.** Most frequent shortest paths terminating at the DHF substrate in the WT DHFR system. The table details specific sequences of residues forming hydrogen-bond paths that end at DHF. Occurrences were calculated across 2000 snapshots from the MD simulations. Only pathways with an occurrence greater than 500 are reported. The data illustrates how distal structural signals frequently funnel through common hub residues, most notably D27 and T113, to reach the substrate.

| WT |  |
| --- | --- |
| Path | Occurrence |
| L4, Y111, A6, **T113**, **D27**, DHF | 620 |
| Y111, A6, **T113**, **D27**, DHF | 670 |
| I115, L8, **T113**, **D27**, DHF | 1170 |
| A6, **T113**, **D27**, DHF | 896 |
| L8, **T113**, **D27**, DHF | 1325 |
| T35, F31, **D27**, DHF | 1011 |
| T113, **D27**, DHF | 1456 |
| L28, K32, DHF | 872 |
| F31, **D27**, DHF | 1227 |
| I50, R52, DHF | 535 |

In addition to the path count above, two complementary measures confirm the sparse, hub-organized nature of WT DHF-directed connectivity. First, the time-averaged path lifetimes (**Figure S5a**) reveal persistent DHF-directed connectivity concentrated at L8, D27, R52, and T113, with R52 exhibiting the highest lifetime peak in the network. The average path lengths (**Figure S5b**) fluctuate broadly with large standard deviations, consistent with the transient character of extended hydrogen-bond routes, so persistence rather than length discriminates the functionally relevant residues. Second, the instantaneous incoming path count (**Figure S6**) quantifies how strongly DHF-directed shortest paths converge on each intermediate residue. Only a limited set of positions displays elevated convergence, most prominently D27 and K58, alongside secondary contributions from T113 and Y111 within the K109–I115 cluster, confirming that pathway flow is channeled through sparse integrators rather than distributed uniformly.

The hub residues identified by these three measures have well-documented biochemical roles. D27 governs the overall rate of DHF-to-tetrahydrofolate conversion and, together with Y100, modulates the pKa of the DHF N5 atom to enable preprotonation [66]. T113V mutagenesis perturbs both ligand binding and catalysis through disruption of the hydrogen-bond network [67], consistent with our identification of T113 as the dominant funneling hub. R57 is critical for substrate-analog binding [68], and T35 is invariant across DHFR homologs and stabilizes the α/β-helix terminus [68]. W22 and N23 lie immediately adjacent to cluster (ii) (L24/D27/F31/K32/T35/K38; see **Figure 3a**) and position the M20 side chain relative to the DHF N5 atom [69]. Through this contact, they link cluster (ii) to the active-site geometry required for catalysis. The convergence of our network-derived hubs on residues with independently characterized catalytic and conformational roles supports the interpretation that DHF-directed connectivity in WT is organized around a small number of integrator nodes.

### Temporal analysis provides a complementary, exploratory view of DHF-directed connectivity in WT

Because ligand-induced allosteric transmission can occur with a time lag, we next asked whether DHF-directed paths are traversed across successive frames on the aggregated graph, introducing latency into the analysis. Two caveats apply throughout this section and should be kept in mind when interpreting the results. First, the total sampling of two 1 µs trajectories (2000 snapshots at 1 ns spacing) captures nanosecond-scale hydrogen-bond dynamics well but is not expected to fully resolve the repetitive formation and disruption of delayed DHF-directed paths, since motions relevant to the DHFR catalytic cycle extend into the millisecond regime [70]. The temporal descriptors reported here should therefore be interpreted with this hierarchy in mind: instantaneous and short-latency descriptors are expected to be the most reliably resolved by our sampling, consistent with the broader picture that allosteric communication is initiated on nanosecond timescales before evolving into the microsecond–millisecond range [35, 36]. Second, the aggregated graph is most meaningful when restricted to stable molecular identities. We therefore treat the non-water aggregated graph as the primary temporal representation.

Extending the incoming path count to the temporal setting (**Figure 4**) suggests that, within the non-water aggregated graph, time-resolved pathway flow converges on a limited set of intermediates, with the strongest peak at N18 and additional contributions from A6, N23, T35, K38, R57, Y100, and Y111. Several of the residues highlighted in **Figure 4** residues have established biochemical roles that align well with this assignment: Y100 participates with D27 in modulating the DHF N5 pKa [66]; T35 and R57 are conformation-stabilizing and ligand-binding residues, respectively [68]; and N23 is positioned at the base of the M20 loop in the active-site region [69]. The convergence between the temporal descriptors and these independently established functional roles supports the interpretation that the delayed-path analysis identifies functionally relevant intermediates.

**Figure 4.**
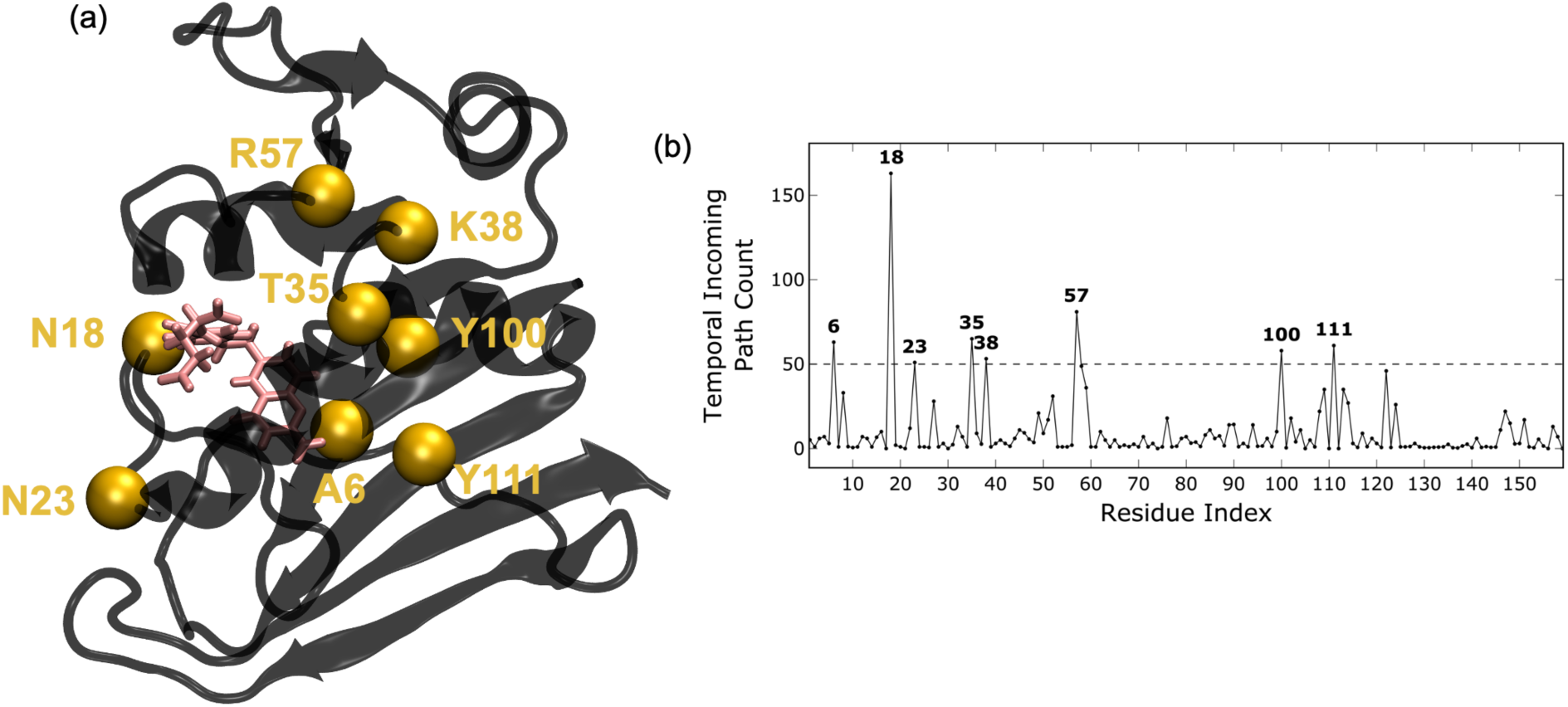
Temporal incoming path count identifies hub-like residues in time-resolved DHF-directed connectivity in WT DHFR. **(a)** Structural representation of WT DHFR highlighting residues with elevated temporal incoming path count as gold spheres (A6, N18, N23, T35, K38, R57, Y100, and Y111). These residues mark intermediate positions that accumulate temporally ordered shortest-path flow toward DHF and therefore act as key integrators of delayed connectivity. DHF and NADPH are shown in stick representation. **(b)** Residue-wise profile of the temporal incoming path count computed from temporal shortest paths on the aggregated graph. Peaks above the dashed threshold identify residues that receive disproportionately many time-resolved paths en route to DHF, revealing the main control points for substrate-directed connectivity when latency is explicitly taken into account.

Taken together, the instantaneous and temporal analyses of WT DHFR provide complementary views of DHF-directed connectivity: a snapshot-wise scaffold funneled through D27 and T113, and a time-ordered layer organized around a distinct group of intermediates that link these endpoints through delayed routes, offering a more complete picture of allosteric communication than either descriptor alone. Longer-timescale simulations are expected to further refine the temporal layer.

### Instantaneous hydrogen-bond paths are conserved in the L28R mutant while gatekeeper residues are shifted

The L28R substitution is expected to perturb connectivity without causing large structural rearrangements, and we therefore applied the same instantaneous and temporal framework to L28R to ask which features of the WT network are preserved.

In L28R, the direct hydrogen-bond interface to DHF is dominated by D27, R28, and R52 (**Table S1**), with occurrences of 1837, 1615, and 1561, respectively. Compared with WT (**Figure 2d**), D27 and R52 are retained as dominant contacts, while the newly introduced R28 side chain establishes a persistent direct interaction with DHF and K32 contributes less frequently than in WT (1140 → 683 snapshots). The interface is therefore largely preserved in its D27/R52 anchoring but locally rewired by partial replacement of K32 contacts with R28 contacts. This localized rewiring is consistent with the established mechanism of L28R-mediated trimethoprim resistance, in which the arginine side chain forms additional hydrogen bonds with the glutamate tail of DHF and thereby increases substrate affinity relative to inhibitor affinity [8, 46, 53].

The instantaneous descriptors are partially conserved between WT and L28R. The total instantaneous shortest-path count per residue directed toward DHF is highly conserved between WT and L28R (*R*^2^ = 0.95, **Figure S7** and **Table S2**), indicating that the overall DHF-directed pathway architecture is largely preserved upon mutation. The residue-wise comparison of the three instantaneous shortest-path descriptors (**Figure S8**) shows that path lifetimes are the most conserved: time-averaged lifetimes correlate strongly between the two systems (*R*^2^ = 0.84, **Figure S8a**), indicating that residues maintaining persistent hydrogen-bond communication with DHF in WT largely retain this feature in the mutant (**Figure S9**). The average shortest-path length and the instantaneous incoming path count show moderate correlations (*R*^2^ = 0.35 and *R*^2^ = 0.42, respectively; **Figure S8b,c**), reflecting snapshot-level rewiring of how individual residues route their paths to DHF. This rewiring is also visible in the mutant incoming path count profile (**Figure S10**), where D27 remains a dominant integrator while additional contributions appear at T46, I50, A127, and Q143 alongside preserved peaks at E90, Y111, T113, and E157.

The temporal descriptors reveal a redistribution of delayed communication in L28R. The residue-wise correlations of temporal descriptors (**Figure S11**) are weaker than for the instantaneous measures, indicating that the general architecture of the aggregated graph is substantially altered upon mutation (**Figure S11a,b**), with temporal incoming path counts essentially decorrelated (*R*^2^ = 0.16, **Figure S11c**). **Figure S12** shows a corresponding redistribution of the temporal incoming path count: while several WT-dominant hubs are diminished (N18, A6, T35, Y100, Y111; **Figure 4**), others are strongly retained and amplified (N23, K38, R57), shifting pathway flow alongside the emergence of new mutant-specific hubs including A7, M20, W22, T46, I50, R52, N59, and T113, with accumulation in the T46–R57 region. This pattern is consistent with the instantaneous analyses and with the known mechanism of L28R [8, 46, 53], in which local rewiring around the mutation site reshapes how delayed routes traverse the network while preserving overall path availability.

The redistributed temporal residues are enriched in positions with well-characterized dynamic and electrostatic relevance to DHFR catalysis. M20 and W22 belong to the active-site machinery that protonates DHF whereby the M20 loop provides the final pKa boost for substrate protonation from solvent [71], and W22 positions the M20 side chain relative to the DHF N5 atom [69]. T46 has been used as a thiocyanate vibrational probe (T46C) to map the heterogeneous electrostatic microenvironment of the active site [72]. The convergence of the temporal redistribution onto these functionally characterized residues supports the interpretation that L28R reroutes delayed communication through positions that are themselves coupled to catalysis.

### Network hotspots partially overlap with coevolution and cryptic-site predictors

To place the network-derived hotspots in a broader evolutionary and structural context, we compared the residue-resolved connectivity profiles with independent cryptic-site and coevolution analyses (**Figure 5**) [19, 21, 24, 30, 31]. The cryptic-site predictors (PocketMiner, CryptoSite, PlayMolecule) do not identify a single dominant hotspot but highlight several discrete peaks along the sequence, indicating that latent pocket-forming propensity is unevenly distributed across the DHFR scaffold. The sequence-based coupling analyses (GREMLIN, EVcouplings) show pronounced enrichment in the C-terminal half of the protein, with the strongest cumulative coupling in residues 100–120.

**Figure 5.**
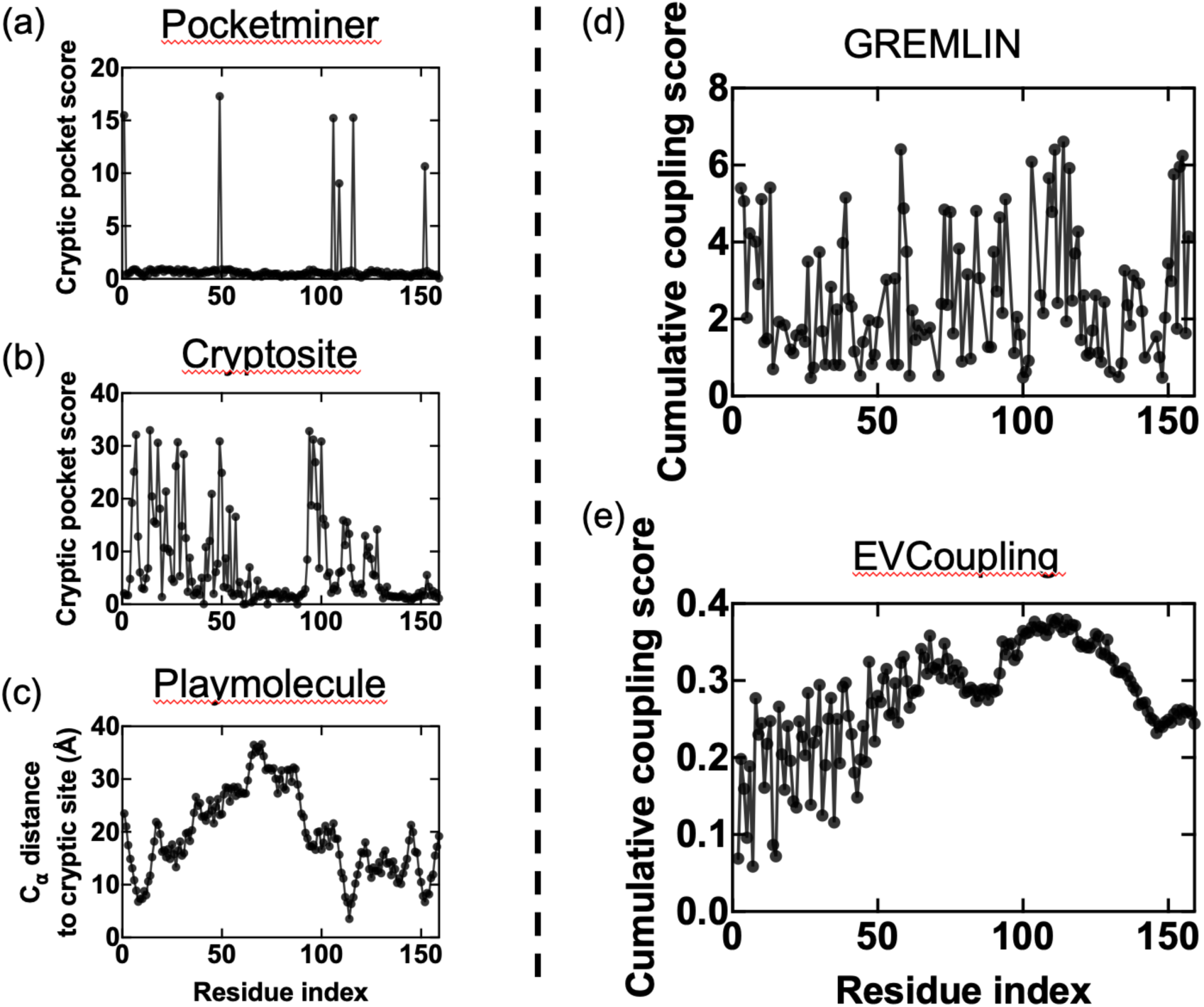
Cryptic-site and coevolution analyses of DHFR. **(a)** PocketMiner cryptic-pocket score as a function of residue index. **(b)** CryptoSite cryptic-pocket score as a function of residue index. **(c)** Distance of each residue C_α_ atom to the cryptic site identified by PlayMolecule. **(d)** Cumulative coupling score obtained from GREMLIN. **(e)** Cumulative coupling score obtained from EVcouplings. Together, these panels provide complementary views of distal functional coupling in DHFR by comparing residues predicted to contribute to cryptic-site formation with residues highlighted by sequence-based coevolution analyses, enabling qualitative comparison with the hydrogen-bond communication hotspots identified in the network-based analyses.

The K109–I115 segment, identified in the instantaneous analyses as a recurrent and persistent component of DHF-directed connectivity, coincides with a prominent peak in both coevolution profiles. Overlap with the cryptic-site predictors is more limited, suggesting that hydrogen-bond network analysis captures features of distal communication that are only partially recovered by static structural approaches. Independent experimental evidence further supports the functional relevance of this segment and of the broader structural context in which it is embedded. First, K109–I115 lies within a secondary allosteric pocket that conformationally couples to the M20 loop [49], so that residues in and around this segment are positioned to mediate long-range coupling to the active site. Second, deep mutational scanning of DHFR has identified a distributed set of advantageous but destabilizing hotspots (including L24, W47, H114, D116, and E154) [73], of which H114 and D116 fall directly within or immediately adjacent to the K109–I115 segment. Third, morbidostat evolution experiments on TMP-resistant DHFR recurrently select mutations at W30, D116, H141, and D144 [8]; D116 again falls within the K109–I115 segment, while the remaining positions cluster in the same allosteric pocket. The convergence, at this structural region, of the hydrogen-bond network, coevolutionary coupling, allosteric-pocket formation, and evolutionarily accessible resistance mutations reinforces the interpretation that DHFR harbors a genuine communication sector centered on K109–I115 and its surrounding pocket, whose functional relevance is supported by multiple independent lines of evidence.

## Conclusion

In summary, we present a hydrogen-bond network framework that integrates instantaneous and temporal graph analyses to quantify ligand-directed connectivity directly from MD trajectories. Applied to *E. coli* DHFR, the method shows that substrate-directed connectivity is not distributed uniformly across the protein but is instead organized through a sparse and dynamically evolving set of preferred routes. In the wild-type enzyme, these routes recurrently converge on a limited number of hub-like residues, with D27 and T113 emerging as particularly important integrators of DHF-directed connectivity.

Comparison with the L28R mutant shows that the mutation neither disrupts the core substrate-contacting interface nor abolishes the overall communication scaffold. Its primary effect is instead to redistribute path usage, persistence, and temporal convergence within a conserved network. The temporal analyses extend this interpretation by showing that delayed communication is itself highly selective, and that the residues controlling latency-constrained pathway flow shift in a mutation-dependent manner. In fact, this shift may be directly related to the redirecting of the evolutionary path of DHFR away from this mutation in the presence of evolutionary pressure from a carefully constructed trimethoprim derivative that manipulates the local dynamics in the binding pocket [46].

Cross-referencing the network results with independent coevolution and cryptic-site analyses reveals partial but informative agreement whereby network-derived hotspots in the K109–I115 region coincide with a prominent coevolution peak, while overlap with cryptic-site predictors is more limited. This pattern suggests that hydrogen-bond network analysis captures features of distal communication that are only partially recovered by static structural or sequence-based approaches. Because the procedure relies only on hydrogen-bond geometries and time-ordered trajectories, the framework is broadly transferable to other ligand-binding proteins for identifying persistent, delayed, and mutation-sensitive signaling pathways that are not readily apparent from static analyses alone. By operating natively on the nanosecond–microsecond window, the framework targets the timescale at which allosteric signals are first established [32, 34], and can be extended to longer-timescale or enhanced-sampling trajectories to capture delayed communication routes that lie beyond the time scales reachable by conventional MD.

## Acknowledgements

The numerical calculations reported in this paper were partially performed at the Scientific and Technological Council of Türkiye National Academic Network and Information Center (TUBITAK ULAKBIM), High Performance and Grid Computing Center (TRUBA resources).

## Data Availability

The Python implementation of the temporal hydrogen-bond network analysis pipeline is openly available on GitHub at https://github.com/tfguclu/temporal_hydrogen_bond_network. The molecular dynamics trajectories and the raw data supporting the findings of this study are openly available on Zenodo at https://doi.org/10.5281/zenodo.20024225.

## Supporting Information

Correspondence between the DHF-directed incoming path count and DHF-directed betweenness centrality in WT DHFR (**Figure S1**); time evolution of the number of edges in the WT DHFR residue network (**Figure S2**); probability distributions of hydrogen-bond lifetimes for protein–protein, protein–water, and water–water interactions (**Figure S3**); probability distributions of shortest paths and their characteristics, including path multiplicity, path length, and consecutive water occupancy (**Figure S4**); average lengths and lifetimes of shortest hydrogen-bond paths to the DHF substrate in WT DHFR (**Figure S5**); instantaneous incoming path count identifying hub-like residues in WT DHFR communication toward DHF (**Figure S6**); occurrences of direct hydrogen bonds between L28R DHFR residues and the DHF substrate (Table S1); residue-resolved shortest paths to DHF in the L28R mutant and their correlation with the WT profile (**Figure S7**); most frequent shortest paths terminating at DHF in the L28R DHFR system (Table S2); residue-wise correlations of instantaneous shortest-path lifetimes, lengths, and incoming path counts between WT and L28R (**Figure S8**); average lengths and lifetimes of shortest hydrogen-bond paths to DHF in L28R DHFR (**Figure S9**); instantaneous incoming path count for the L28R mutant (**Figure S10**); residue-wise correlations of temporal shortest-path properties between WT and L28R in the non-water aggregated representation (**Figure S11**); temporal incoming path count revealing mutant-specific hub residues in L28R DHFR (**Figure S12**).

## TOC graphic

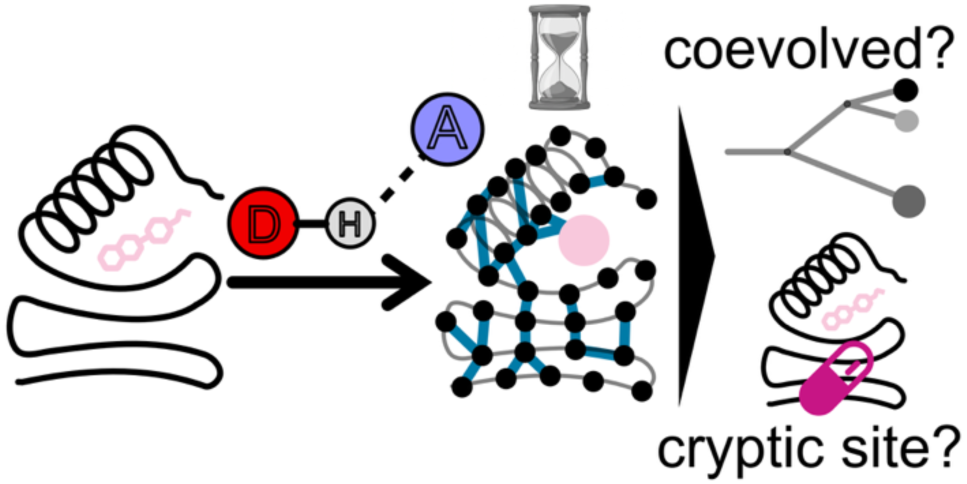

## Supporting Information for

**Figure S1.**
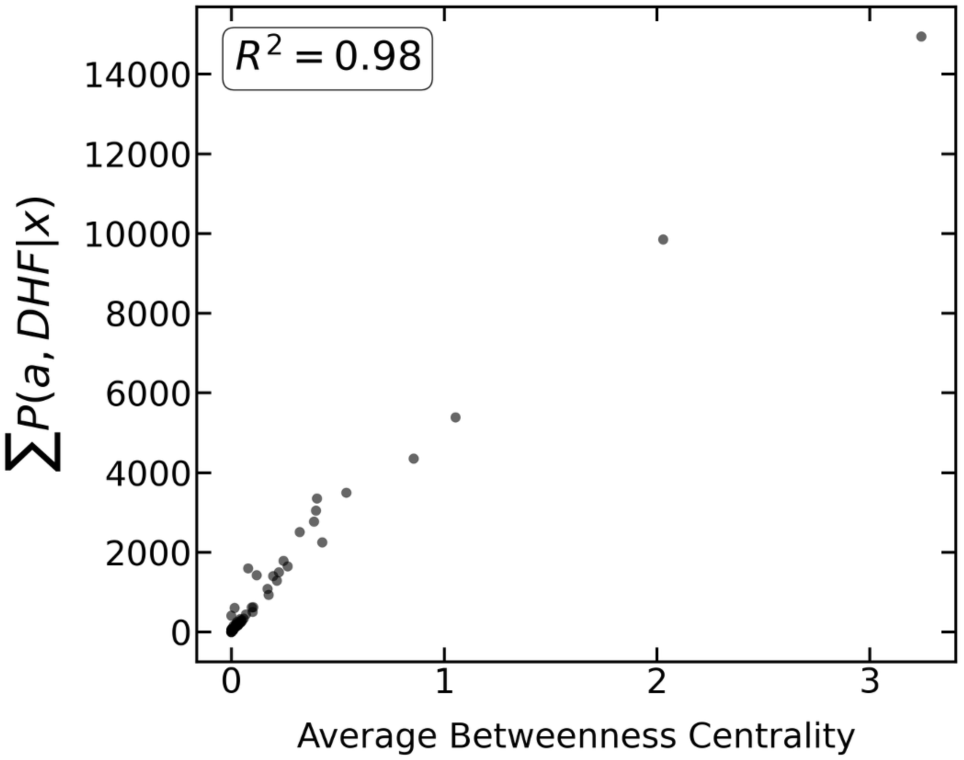
Strong correspondence between the DHF-directed Incoming Path Count and DHF-directed betweenness centrality in WT DHFR. Scatter plot of residue-wise ∑ *P*(*a*, *DHF*|*x*) against time-averaged betweenness centrality computed from the instantaneous hydrogen-bond graphs with DHF as the sink. Each point represents a residue. The high coefficient of determination (*R*^2^ = 0.98) indicates that the Incoming Path Count closely tracks DHF-directed betweenness centrality while providing a computationally less expensive description of substrate-directed pathway convergence.

**Figure S2.**
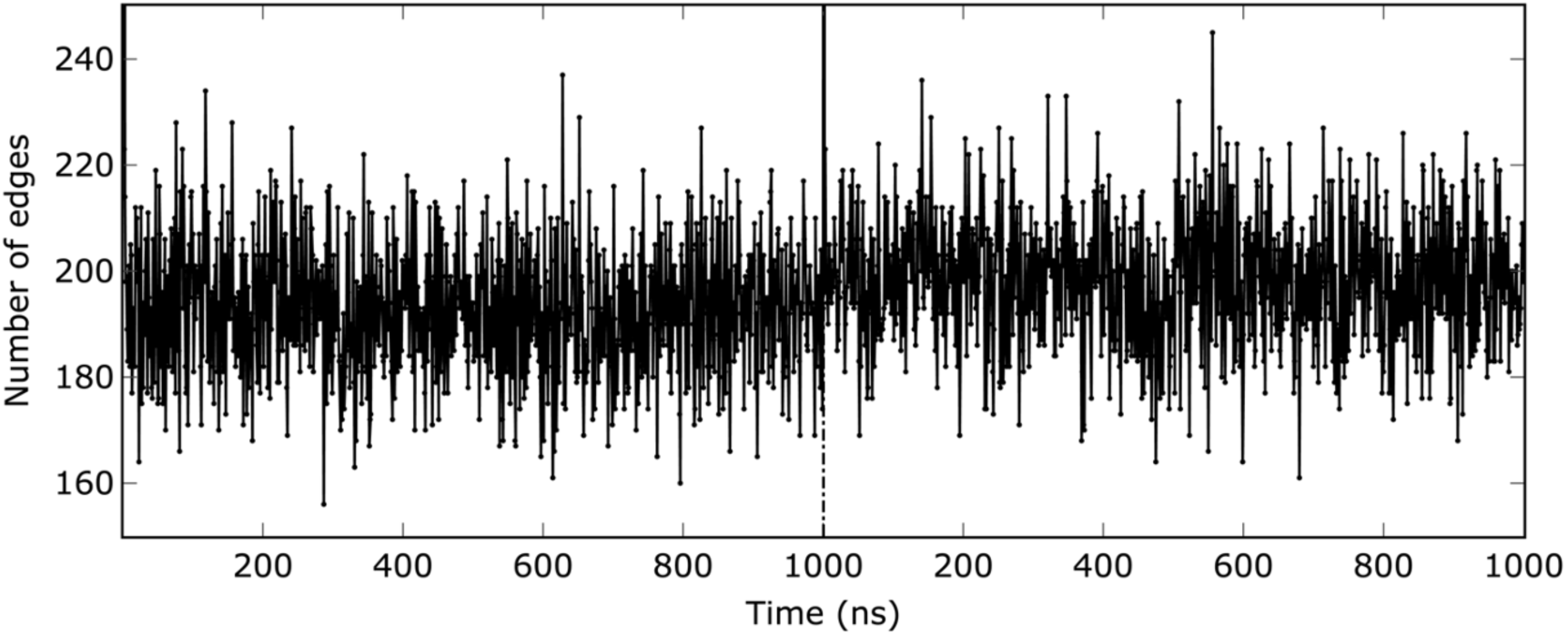
Time evolution of the number of edges in the wild-type (WT) DHFR residue network. The plot displays the total number of edges (representing hydrogen bonds) across two independent 1 µs molecular dynamics (MD) trajectories. The two trajectories are separated by the vertical dashed line. Throughout the simulations, the number of edges fluctuates around a consistent average of approximately 200, excluding the initial dense snapshot of each trajectory.

**Figure S3.**
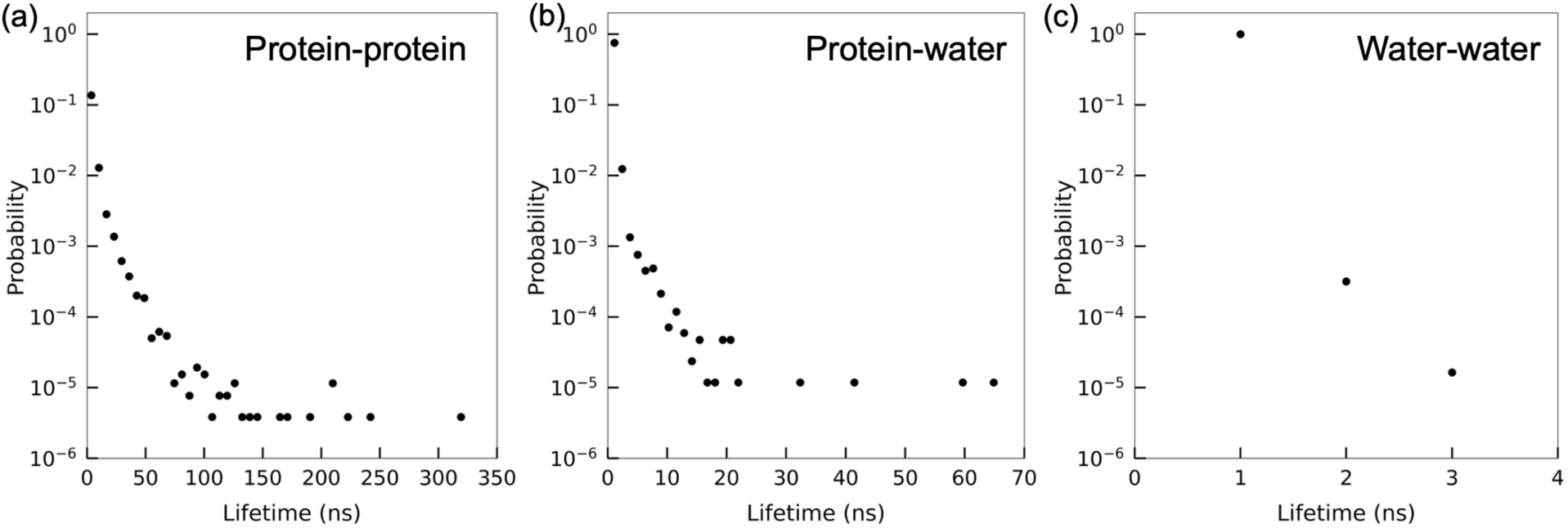
Probability distributions of hydrogen bond lifetimes in the DHFR system. The scatter plots display the lifetime distributions. Protein–protein interactions **(a)** exhibit the longest lifetimes (extending up to ∼300 ns) due to internal structural constraints. Protein–water bonds **(b)** show intermediate lifetimes, while water–water interactions **(c)** are highly dynamic and possess the shortest lifetimes, predominantly around 1 ns. The *y*-axis (Probability) is shown on a logarithmic scale.

**Figure S4.**
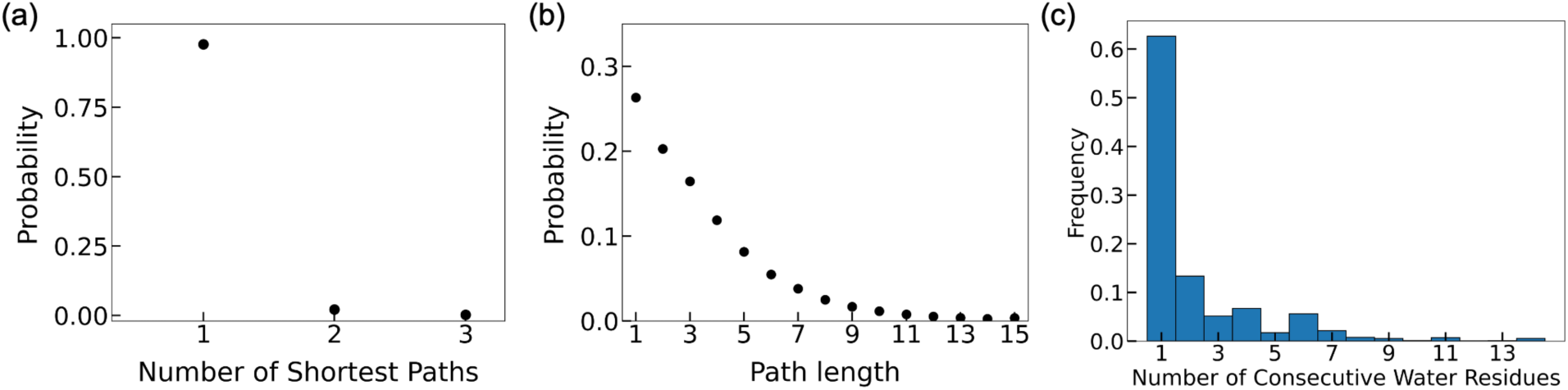
Probability distributions of shortest paths and their characteristics in the DHFR system. **(a)** The probability distribution for the number of shortest paths between any two connected nodes, showing that connected nodes share exactly one shortest path in the vast majority of cases. **(b)** Probability distribution of shortest-path lengths between connected node pairs. Direct contacts (length 1) are the most probable, with the probability decaying monotonically up to approximately 15 edges. **(c)** The frequency distribution of the number of consecutive water nodes within these pathways. When water molecules are present in a path, they most commonly appear as single bridges or in short chains of 2 - 4 molecules, though extended chains can occasionally occur.

**Figure S5.**
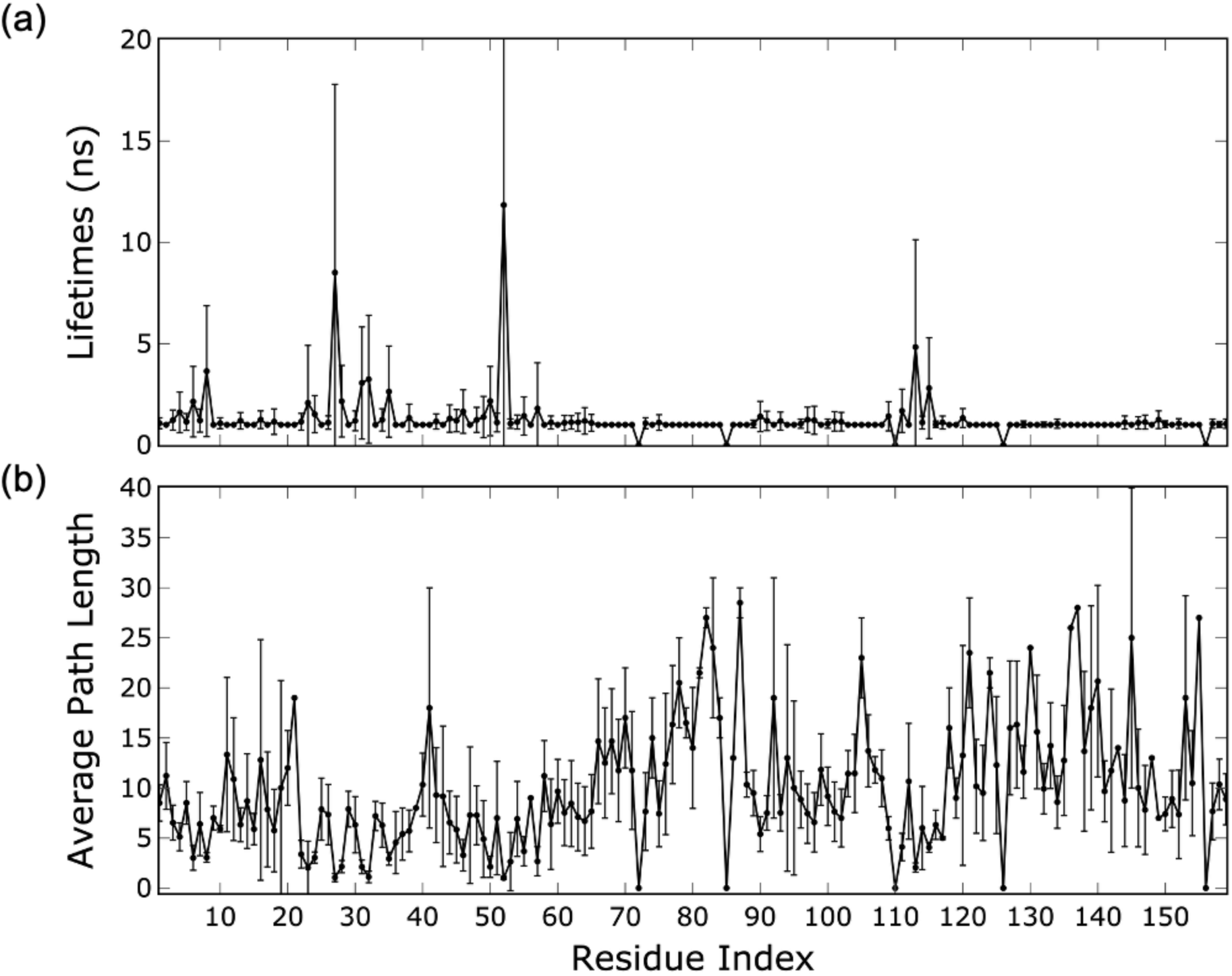
Average lengths and lifetimes of shortest hydrogen-bond paths to the DHF substrate in the WT system. **(a)** The average lifetimes of these shortest paths, defined by their consecutive occurrences in nanoseconds. Distinct peaks reveal residues that maintain highly persistent communication with DHF, notably highlighting key anchoring and hub residues such as L8, D27, R52, and T113. **(b)** The time-averaged path length (number of edges) from each protein residue to DHF. The error bars represent the standard deviation across the simulation snapshots, illustrating the structural fluctuations and dynamic nature of these communication routes.

**Figure S6.**
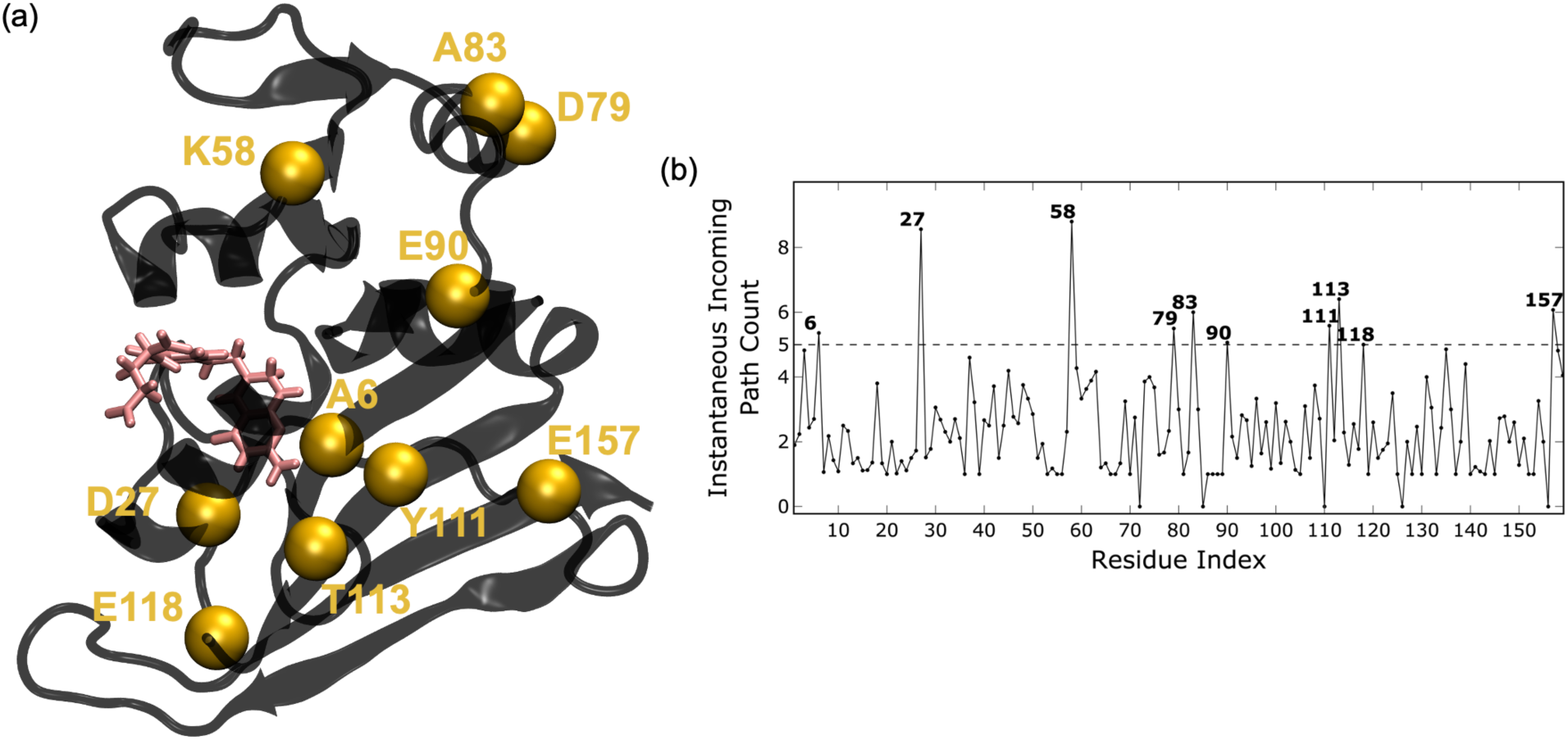
Instantaneous incoming path count identifies hub-like residues in WT DHFR communication toward DHF. **(a)** Structural representation of DHFR showing residues with elevated instantaneous incoming path count as gold spheres (A6, D27, K58, D79, A83, E90, Y111, T113, E118, and E157); DHF is shown in stick representation. These residues mark intermediate positions that collect shortest hydrogen-bond communication pathways directed toward the substrate. **(b)** Residue-wise profile of the instantaneous incoming path count. Peaks above the dashed threshold highlight residues that receive disproportionately many shortest paths *en route* to DHF, revealing a sparse set of funneling/integrating hubs distributed across the protein scaffold. This interpretation follows our definition of incoming path count as a target-directed measure of convergent pathway flow toward DHF.

**Table S1.** Occurrences of direct hydrogen bonds between L28R DHFR residues and the DHF substrate. The occurrences represent the number of snapshots in which a direct hydrogen bond was detected between the specified protein residue and DHF across 2000 frames from two 1 µs MD simulations of the L28R mutant.

| L28R |  |
| --- | --- |
| Residue | Occurrence |
| M20 | 8 |
| N23 | 6 |
| D27 | 1837 |
| R28 | 1615 |
| K32 | 683 |
| R52 | 1561 |
| R57 | 281 |
| T113 | 34 |

**Figure S7.**
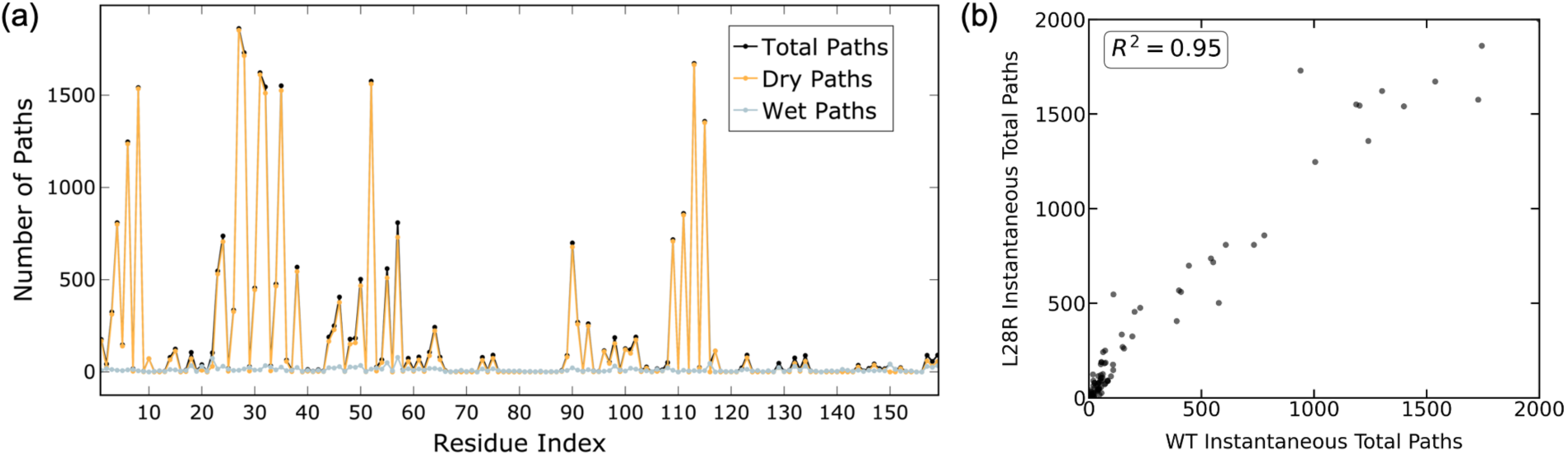
Hydrogen-bond shortest paths directed toward the DHF substrate in the L28R mutant and their correspondence with the wild-type (WT) profile. **(a)** The total count of shortest paths originating from each residue and terminating at DHF, computed across 2000 MD snapshots from two 1 µs simulations of L28R. Pathways are separated into dry paths (excluding water nodes), wet paths (including water nodes), and total paths, following the instantaneous shortest-path analysis used throughout the manuscript. **(b)** Correlation between the WT and L28R instantaneous total-path counts for each residue. The strong agreement between the two profiles (*R*^2^ = 0.95) indicates that the overall DHF-directed pathway architecture is largely conserved between WT and mutant.

**Table S2.** Most frequent shortest paths terminating at the DHF substrate in the L28R DHFR system. The table lists the most frequently observed hydrogen-bond paths that end at DHF in the L28R mutant, with occurrences accumulated over 2000 snapshots from two 1 µs MD simulations. Only pathways with an occurrence greater than 500 are reported. As in the WT system, the most recurrent paths in L28R largely funnel through the D27/T113 axis, while an additional prominent branch involving residue 28 is also retained, consistent with conservation of the core DHF-directed pathway architecture despite mutational rewiring.

| L28R |  |
| --- | --- |
| Path | Occurrence |
| L4, Y111, A6, **T113**, **D27**, DHF | 694 |
| Y111, A6, **T113**, **D27**, DHF | 754 |
| I115, L8, **T113**, **D27**, DHF | 1312 |
| A6, **T113**, **D27**, DHF | 1184 |
| L8, **T113**, **D27**, DHF | 1490 |
| T35, F31, **D27**, DHF | 1411 |
| **T113**, **D27**, DHF | 1615 |
| K32, R28, DHF | 820 |
| F31, **D27**, DHF | 1582 |
| L24, R28, DHF | 654 |

**Figure S8.**
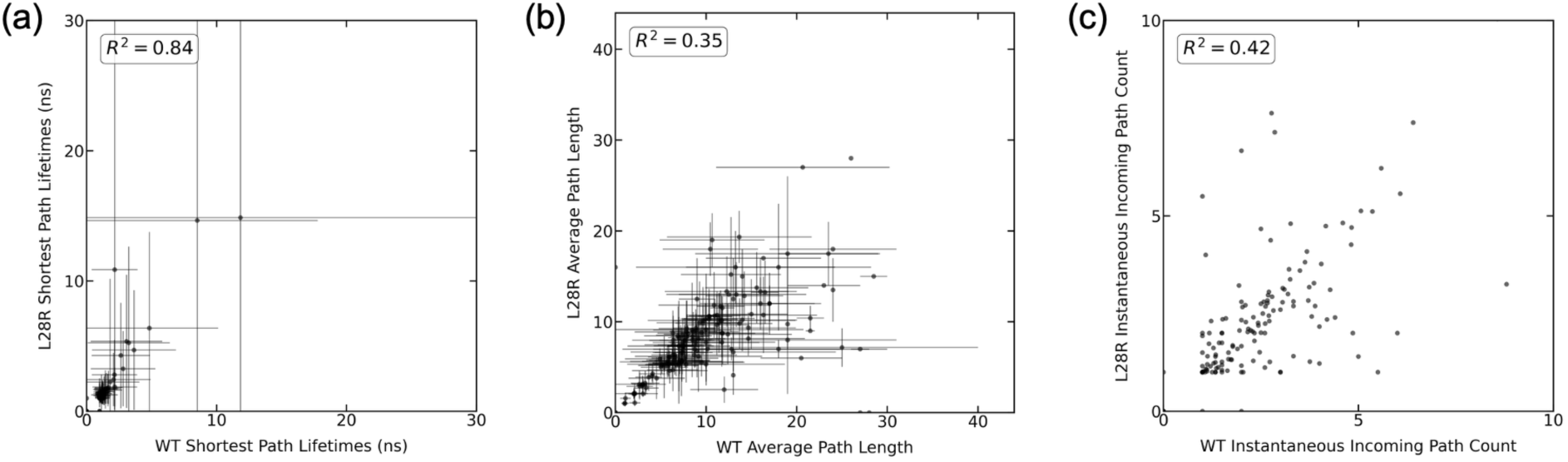
Residue-wise comparison of instantaneous shortest-path properties between wild-type (WT) and L28R DHFR. **(a)** Correlation of average shortest-path lifetimes to DHF between WT and L28R. **(b)** Correlation of average shortest-path lengths between WT and L28R. **(c)** Correlation of the instantaneous incoming path count between WT and L28R. The coefficients of determination (*R*^2^) indicate that shortest-path lifetimes are strongly correlated between WT and mutant, whereas average path length and instantaneous incoming path count show only limited correlation, consistent with preservation of the overall DHF-directed communication scaffold together with mutation-dependent redistribution of pathway usage and hub preference.

**Figure S9.**
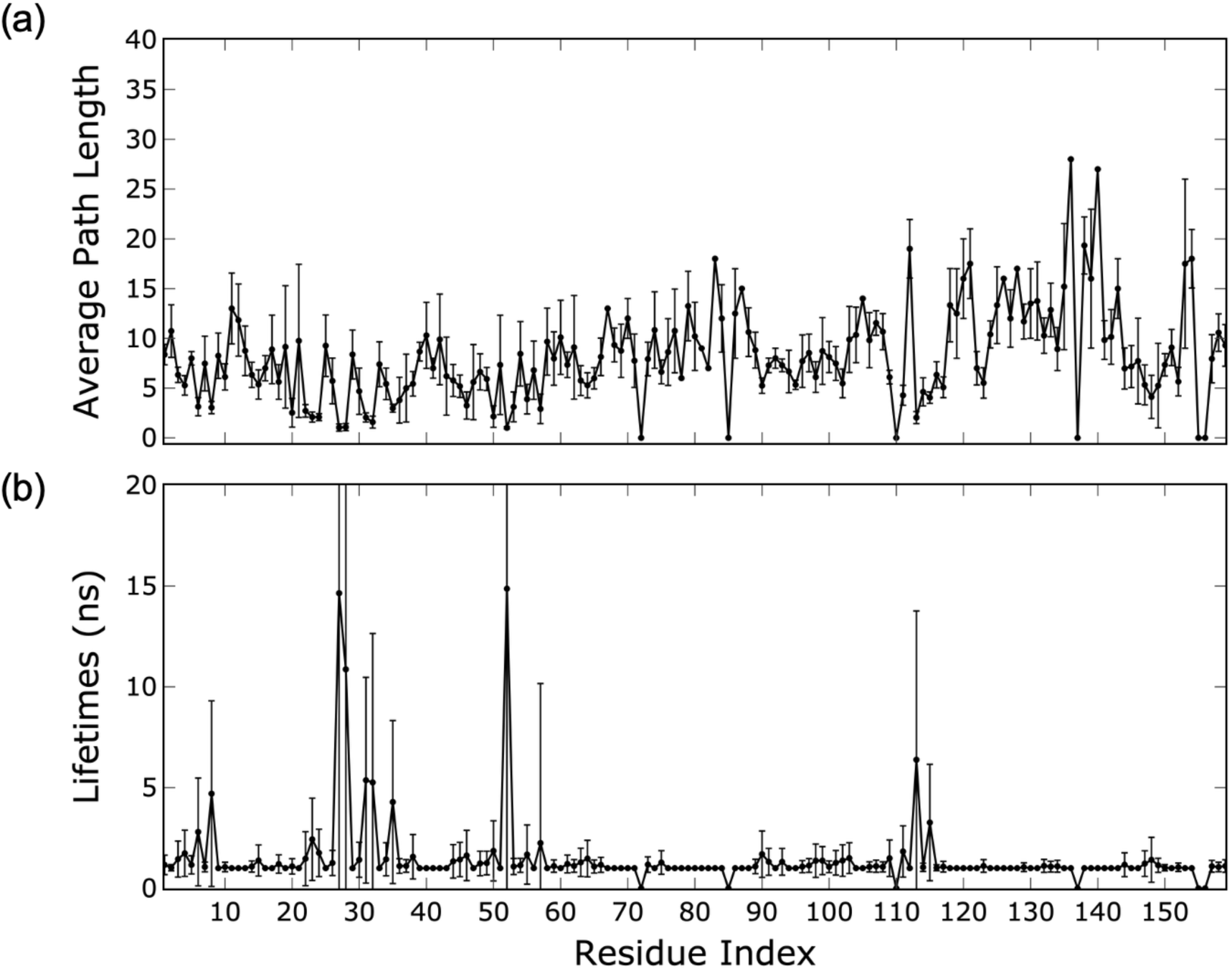
Average lengths and lifetimes of shortest hydrogen-bond paths to the DHF substrate in the L28R DHFR system. **(a)** Time-averaged shortest-path length from each protein residue to DHF, reported as the number of edges; error bars represent the standard deviation across the simulation snapshots. **(b)** Average lifetimes of these shortest paths, defined from consecutive occurrences and reported in nanoseconds (ns). The analysis was performed for the L28R mutant using the same instantaneous shortest-path framework applied to WT across 2000 snapshots from two 1 µs MD simulations, providing a residue-resolved view of how the mutation redistributes path persistence within the conserved DHF-directed communication network.

**Figure S10.**
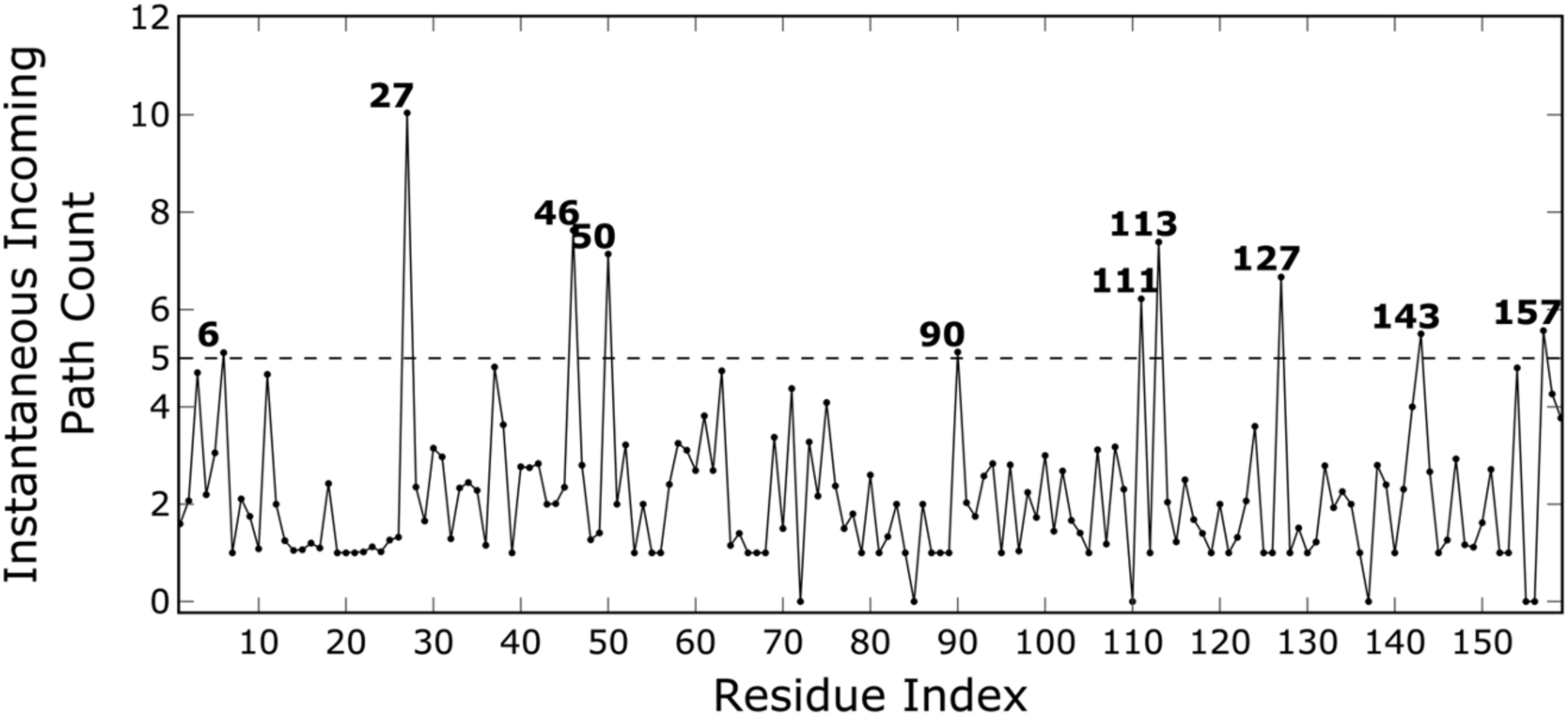
Instantaneous incoming path count identifies hub-like residues in L28R DHFR communication toward DHF. The plot shows the residue-wise profile of the instantaneous incoming path count computed from DHF-directed shortest hydrogen-bond communication pathways in the L28R mutant. Peaks above the dashed threshold highlight residues that receive disproportionately many shortest paths en route to the substrate, revealing a sparse set of funneling/integrating hubs distributed across the protein scaffold. Compared with the WT profile in Figure S6, the mutant retains prominent contributions from residues such as D27, E90, Y111, T113, and E157, while also redistributing pathway convergence toward positions including T46, I50, A127, and Q143.

**Figure S11.**
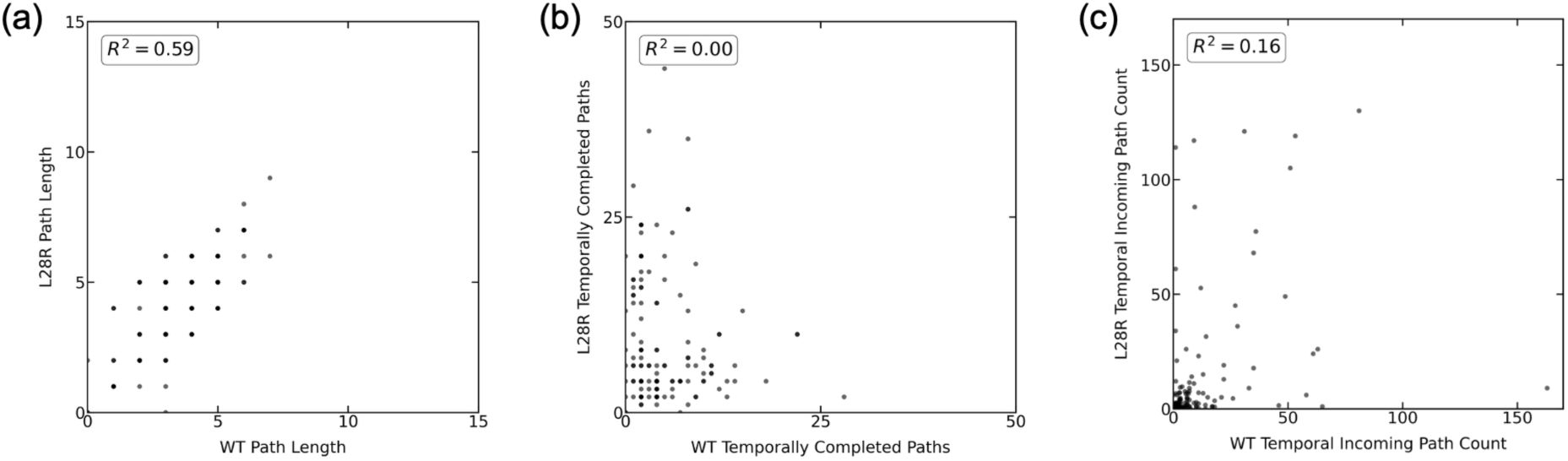
Residue-wise comparison of temporal shortest-path properties between wild-type (WT) and L28R DHFR in the reduced, non-water representation. **(a)** Correlation of aggregated-graph path lengths between WT and L28R. **(b)** Correlation of the number of temporally completed shortest paths between WT and L28R. **(c)** Correlation of the corresponding temporal incoming path count between WT and L28R. The coefficients of determination (*R*^2^) indicate moderate correlation for path length, whereas no correlation is observed for temporally completed paths or for the corresponding temporal incoming path count, consistent with preservation of only the broad temporal communication framework together with strong mutation-dependent redistribution of delayed DHF-directed connectivity.

**Figure S12.**
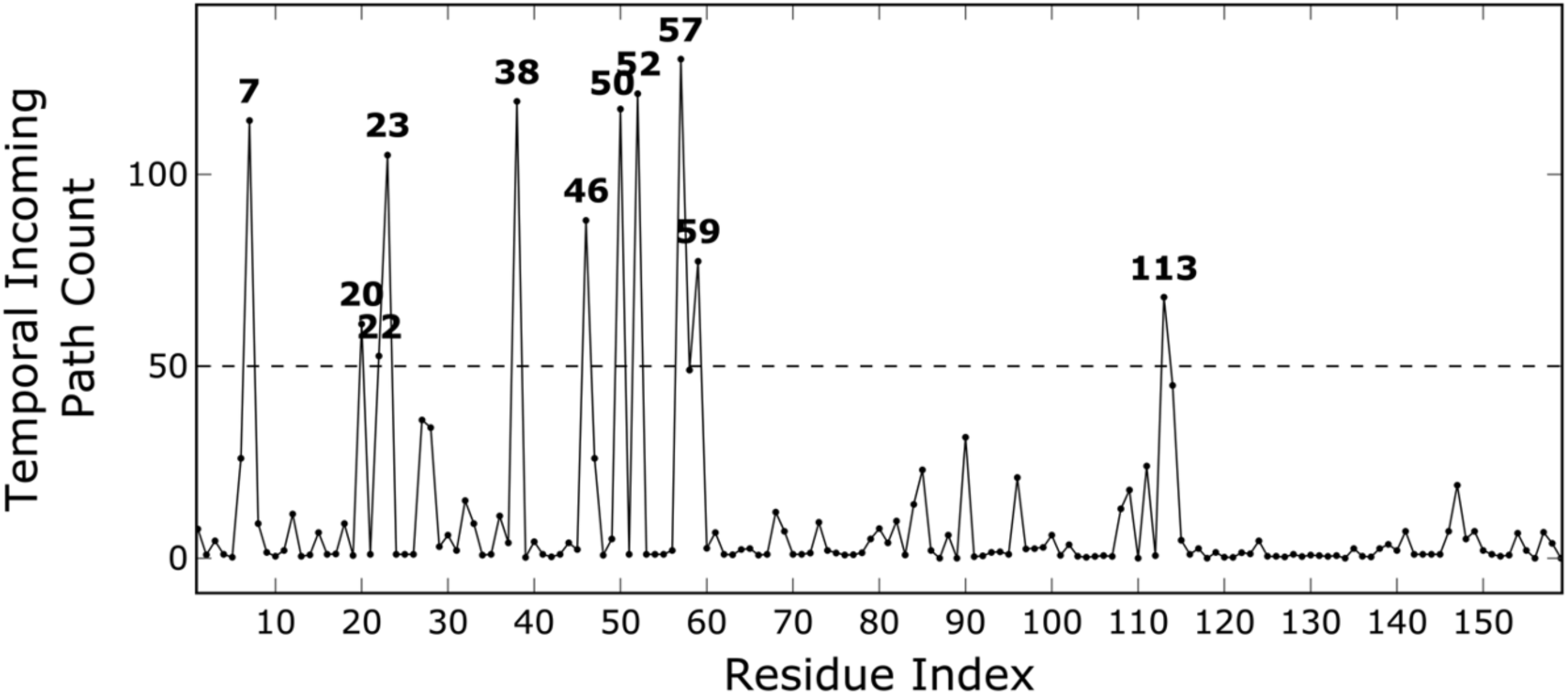
Temporal incoming path count reveals mutant-specific hub residues in time-resolved DHF-directed communication in L28R DHFR. The plot shows the residue-wise profile of the temporal incoming path count computed from temporal shortest paths on the aggregated graph. Peaks above the dashed threshold identify residues that receive disproportionately many time-resolved paths *en route* to DHF, highlighting a sparse set of intermediate positions that act as key integrators of delayed communication in the mutant. Compared with the WT, in which temporal pathway convergence is concentrated at a distinct set of hubs centered on N18 (Figure 4), the L28R profile indicates a redistribution of latency-dependent communication toward residues A7, M20, W22, N23, K38, T46, I50, R52, R57, N59, and T113.

